# LanceOtron: a deep learning peak caller for ATAC-seq, ChIP-seq, and DNase-seq

**DOI:** 10.1101/2021.01.25.428108

**Authors:** Lance D. Hentges, Martin J. Sergeant, Damien J. Downes, Jim R. Hughes, Stephen Taylor

## Abstract

ATAC-seq, ChIP-seq, and DNase-seq have revolutionized molecular biology by allowing researchers to identify important DNA-encoded elements genome-wide. Regions where these elements are found appear as peaks in the analog signal of an assay’s coverage track, and despite the ease with which humans can visually categorize these regions, meaningful peak calls from whole genome datasets require complex analytical techniques. Current methods focus on statistical tests to classify peaks, reducing the information-dense peak shapes to simply maximum height, and discounting that background signals do not completely follow any known probability distribution for significance testing. Deep learning has been shown to be highly accurate for image recognition, on par or exceeding human ability, providing an opportunity to reimagine and improve peak calling. We present the peak calling framework LanceOtron, which combines multifaceted enrichment measurements with deep learning image recognition techniques for assessing peak shape. In benchmarking transcription factor binding, chromatin modification, and open chromatin datasets, LanceOtron outperforms the long-standing, gold-standard peak caller MACS2 through its improved selectivity and near perfect sensitivity. In addition to command line accessibility, a graphical web application was designed to give any researcher the ability to generate optimal peak calls and interactive visualizations in a single step.

## Introduction

Gene regulation is central to cell type specific function and identity, and is dysregulated in disease. Understanding the genomic basis of gene regulation requires mapping regions of protein binding or chromatin modification using methods such as ChIP-seq. Similarly, identifying active regions, as detected by altered chromatin accessibility using ATAC-seq or DNase-seq, provides cell type specific maps of functional regions in the genome. Integrated data from these assays form the high-resolution maps for the main types of genomic elements (enhancers, promoters, and boundary elements) which dictate gene expression in a cell type specific manner^1^. Therefore, the accurate extraction of biologically meaningful data from such assays provides the foundations of current functional genomics research and is critical to understanding gene regulation in health and disease.

Data from ATAC-seq, ChIP-seq, and DNase-seq are processed in a similar fashion: enriched DNA fragments are sequenced, aligned to the genome, and areas enriched for these fragments are recorded. These data appear as tracks of analog signal across genomic coordinates, and increases in fragment density at true-positive biological events are called “peaks'' because of the characteristic pattern of fragments produced in these areas. Besides these regions, enrichment also occurs due to biases and noise in the experimental procedures^2^ or systematic mapping errors common to areas of low complexity^3^. Creating algorithms that can distinguish peaks from such experimental and computational noise, and are robust across methodologies, sequencing depth, diverse tissue types, and chromosomal structure has remained a challenge.

Traditionally, real peaks are distinguished from noise using statistical tests that compare enrichment from the region to background, which is assumed to consist of signal generated randomly. While the Poisson distribution models this better than other distributions^4^, background is in fact nonrandom^5^, appearing at increased levels in areas of open chromatin^6^, at sites with inherent sequence bias and over regions of varying copy number^7^. This must be considered when reviewing significance from statistical peak callers, as misclassification will occur at a higher rate than the p-value suggests. Relying solely on these significance scores may lead to high false positive rates, but also leaves room for potential false negatives, with the ratios of these errors depending on the parameters selected. Exacerbating this default settings are routinely used, reducing accuracy nearly 10% on average from tuned parameters when using statistical peak callers^8^. With these tools, errors may be reduced by using matched negative controls (also known as “input tracks”) to calculate the level of background noise, though this increases the time and costs of the experiment. While peak callers such as MACS2^9^ do not strictly require negative control tracks, forgoing them may sacrifice performance^10^. Input tracks do control for some experimental bias but are still sensitive to chromatin activity, making statistical tests more prone to false negatives^6^.

To address the well-known problems of peak callers, analysis pipelines employing quality control steps are common. The Encyclopedia of DNA Elements (ENCODE) consortium hosts numerous chromatin profiling assay datasets^11,12^ and has developed a robust set of guidelines including recommendations for input controls, sequencing depth, library complexity, and exclusion list regions where mapping errors are more prone to occur^13^.

Multiple replicates are encouraged, and procedures exist for combining peak calls for the most efficient reduction in error^14^. Although these extensive measures greatly improve the reproducibility of peak calls, high-throughput visual inspection showed numerous erroneous peak calls remain^15^.

The inability to reproduce published results is a prevalent concern amongst researchers and is due in part to the unintentional misuse of statistics^16^. These issues include overstating the meaningfulness of the statistical test results and conflating significance with effect^17^ - traps commonly used peak callers fall prey to. Peak callers relying on statistical models simplify the complex analog signal of a region into a single value (maximum height) and use it to calculate a p-value. Enrichment is calculated against a background signal incorrectly assumed to follow a known distribution, and this sole measurement is then falsely equated with peak quality and used to filter results. Quality control is typically limited to uploading the significant regions and coverage track to a genome browser such as UCSC^18^ or IGV^19^, where sections of the genome can be manually scanned. Using only these tools makes anything beyond a cursory inspection tedious and impractical, but because of the incomplete link between statistical test results and peak quality, thoroughly exploring and refining peak calls is of particular importance.

Though extremely time consuming when done at scale, researchers have been shown to effectively judge the quality of peaks using a genome browser. Rye et al. measured peak caller performance by creating a dataset of visually verified peak calls using the UCSC genome browser, and inadvertently measured the performance of the humans in the process^20^. They found that transcription factor motifs, known to be associated with true biological signals, were recovered more often from the manually labeled peaks than from the peak callers. Amazingly they also found that 80% of the software’s false positives could be detected even without an input track, because the human peak callers could identify that these regions “lacked the expected visual appearance of a typical ChIP-seq peak”. Furthermore, while classifying regions by eye is seemingly dependent on an individual, Hocking et al. demonstrated a high consistency across labelers when judging peaks^8^. Visual inspection can be a credible method for peak calling, though to do so comprehensively for an entire human genome would be nearly impossible.

Convolutional neural networks (CNNs), a class of deep learning algorithms, have been extremely successful in a number of general pattern detection tasks such as voice recognition and image classification^21^. Indeed, error rates as low as 3.6% have been achieved for image classification^22^, even surpassing the human error rate of 5.1% for the same dataset^23^. These techniques are being applied in biology as well, especially in genomics where there is an overabundance of data available for training and analysis^24^. Tools such as DeepSea^25^ and Bassett^26^ take genomic sequences as input and can predict regulatory genomic features with high accuracy. Proof of principle studies have also shown promise for applying these techniques to peak calling^8,27^.

Here we present LanceOtron, a peak caller utilising deep learning and packaged with a graphical user interface for integrated quality control. LanceOtron improves upon current tools by calculating a multitude of enrichment metrics for each region being assessed and combines these with a CNN trained to recognise the characteristic shape of peaks. This model is designed for open chromatin, transcription factor and chromatin modification ChIP-seq data, and achieves both high sensitivity and selectivity. Our user-friendly webtool has comprehensive filtering capabilities, built-in genome browser, and automatically generated interactive charts. LanceOtron is freely available at https://LanceOtron.molbiol.ox.ac.uk/.

## Results

### LanceOtron: a deep learning based peak caller with embedded visualization tools

The core of LanceOtron’s peak scoring algorithm is a customized deep neural network strategically combined with local enrichment measurements. These enrichment measurements are taken from the maximum number of overlapping reads in a peak compared to its surroundings - chromosome-wide as well as 10 kilobases (kb) to 100 kb regions in 10 kb increments. The measurements are then used in a logistic regression model, which produces an enrichment score. A base pair resolution view of the signal over a 2 kb window, centered on each peak, is then encoded and input into LanceOtron’s CNN. The CNN uses the relationship between the number of overlapping reads at all 2,000 points, i.e., the shape, to determine if the region is a peak arising from a biological event or noise. Finally, a multilayer perceptron combines the outputs from CNN and logistic regression model, as well as the 11 local enrichment measurements to produce an overall peak quality metric called Peak Score (**Fig. 1a**). As this is a supervised machine learning algorithm, training data is required to provide examples of the shapes and enrichments for the peak and noise regions. For this we used 50 datasets from open chromatin, transcription factor and chromatin modification ChIP-seq experiments (**Supplementary Table 1**) reaching a total of 736,753 labeled regions covering 499 Mb of genome (Methods).

**Fig. 1.**
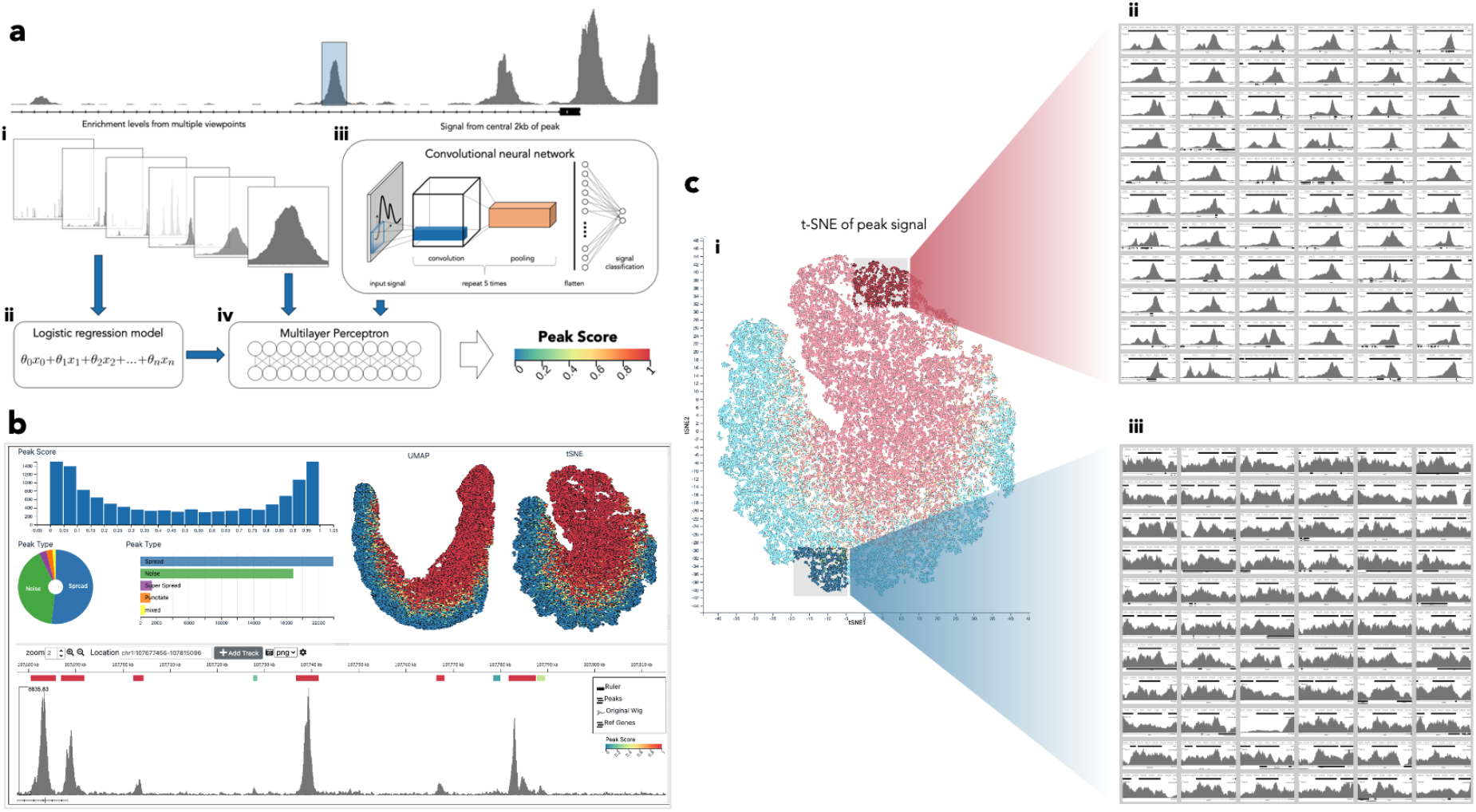
LanceOtron, a deep learning based peak caller overview. **a**, Overview of LanceOtron’s neural network. **i**, Local enrichments are calculated against background from 10kb to 100kb regions in 10kb increments, plus whole chromosome. **ii**, The enrichment values are used as inputs for a logistic regression model. **iii**, Siganl from the central 2 kilobases (kb) is fed into a convolutional neural network (CNN). **iv**, The output from the CNN, logistic regression model, and local enrichment values are all input into a multilayer perceptron, which produces the overall peak score for a given region. **b**, Peak calls are visualized with interactive bed file, charts, clustering, and linked genome browser. Filtering can be applied using LanceOtron’s peak score, p-value, height, genomic coordinates, or any other criteria based on column in the intractive bed file. **c**, Peak call retrieved from ENCODE, but scored with LanceOtron’s model. **i**, Peak calls are clustered and visualized using LanceOtron interactive t-SNE plot, with screen captures from the image thumbnail panel ii, high and iii, low scoring regions as assessed by LanceOtron’s neural network.

LanceOtron extracts genomic data from a bigwig track, which has the benefits of being both compact and readily visualized. With widely used peak callers such as MACS2, assessing the quality of results cannot be done directly, rather the user must upload their output to a genome browser. This is somewhat restrictive for judging the quality of a peak call, in that the output file and genome browser are disconnected, meaning users are limited to haphazardly scanning some genomic regions to see if their results are sensible. To address this LanceOtron is built on the powerful MLV genome visualization software^15^, which allows users to sort and filter results, as well as visualizing peaks and their metadata *en masse*. Clustering peaks based on shape and quality is built-in via the unsupervised machine learning techniques PCA^28^, t-SNE^29^, and UMAP^30^. This allows for rapid assessment of data quality, structure and the appropriateness of the output of the algorithm for the current dataset (**Fig. 1b, Supplementary video 1 & 2**).

LanceOtron’s has three main modules, each taking a coverage file as input and returning enriched regions with associated scores as output. 1) Find and Score Peaks, which first labels enriched regions as candidate peaks, then scores them using LanceOtron’s deep learning model 2) Find and Score Peaks with Inputs performs the same function as the first module but additionally calculates the p-values of regions based on enrichment compared to a separate input control track 3) Score Peaks, which does not find candidate peaks, but rather the neural network scores genomic locations provided as an additional file.

The first two modules, Find and Score Peaks and Find and Score Peaks with Inputs, employ LanceOtron’s candidate peak calling algorithm. This works by applying a 25-way enrichment test, consisting of different smoothing window-threshold combinations (Methods). This allows for various ways for a region to be considered enriched, with the aim of generating an overcomplete set of all possible areas of interest to present to the neural network for assessment. The ethos of LanceOtron is different from existing peak callers, which include or exclude peaks from the output based on parameters and cut-offs. LanceOtron’s aim is to identify all potentially enriched regions, score these using machine learning and return the complete dataset in a manner that can be examined and queried in its entirety. This is made feasible through the comprehensive filtering and powerful data exploration tools LanceOtron’s graphical interface offers. Furthermore, by calculating the comprehensive Peak Score, p-values can be used at relaxed thresholds as a means of excluding peaks found in the input tracks, rather than as the sole means for judging peak quality.

The final module, Score Peaks, uses LanceOtron’s neural network component in isolation from the candidate peak identifier. This allows users to analyze the quality of peak calls from other tools, publications or databases. Using this reanalysis capability, we have found that publicly available peak calls, even following the strictest guidelines, may contain large numbers of low-quality peaks. For example, LanceOtron was used to reanalyze peaks calls from ENCODE ChIP-seq for H3K27ac from 22Rv1 prostate cancer epithelial cells (ENCSR391NPE). As part of the ENCODE pipeline, two biological replicates were independently peak called and only peaks present in both were included. Using LanceOtron’s deep learning based scoring, clustering, and visualization tools it is clear that many very low quality peaks remain in the datasets despite requiring independent calls (LanceOtron 22Rv1 H3K27ac project)(**Fig. 1c**). Large amounts of similarly low-quality peaks can be identified in many other public data sets based on similar statistical peak calling approaches.

### Benchmarking LanceOtron

We benchmarked LanceOtron’s performance with the ENCODE recommended peak caller MACS2, both using default settings (with and without an input control track when available). We compared peak calls from transcription factor ChIP-seq, histone ChIP-seq, and the open chromatin assays ATAC-seq and DNase-seq. For a complete numerical listing of performance benchmarks for all labeled datasets see **Supplementary Table 2**.

#### Transcription factor ChIP-seq

Our transcription factor dataset was CTCF in spleen primary cells, downloaded from ENCODE (ENCSR692ILH). We hand labeled 10 megabases (Mb) of the dataset, marking areas which were obviously peaks or noise (Methods) resulting in 109 human curated peak annotations. When no input control track was used, both LanceOtron and MACS2 achieved perfect sensitivity, detecting all labeled peaks in the dataset, but MACS2 had far lower selectivity and overall F1 score. With input, LanceOtron outperformed MACS2 in precision, recall/sensitivity, selectivity, and F1 score. Comparing across peak call types, LanceOtron without input actually achieved higher scores than MACS2 with input across all metrics (LanceOtron spleen CTCF projects: without input; with input)(**Fig. 2a**).

**Fig. 2.**
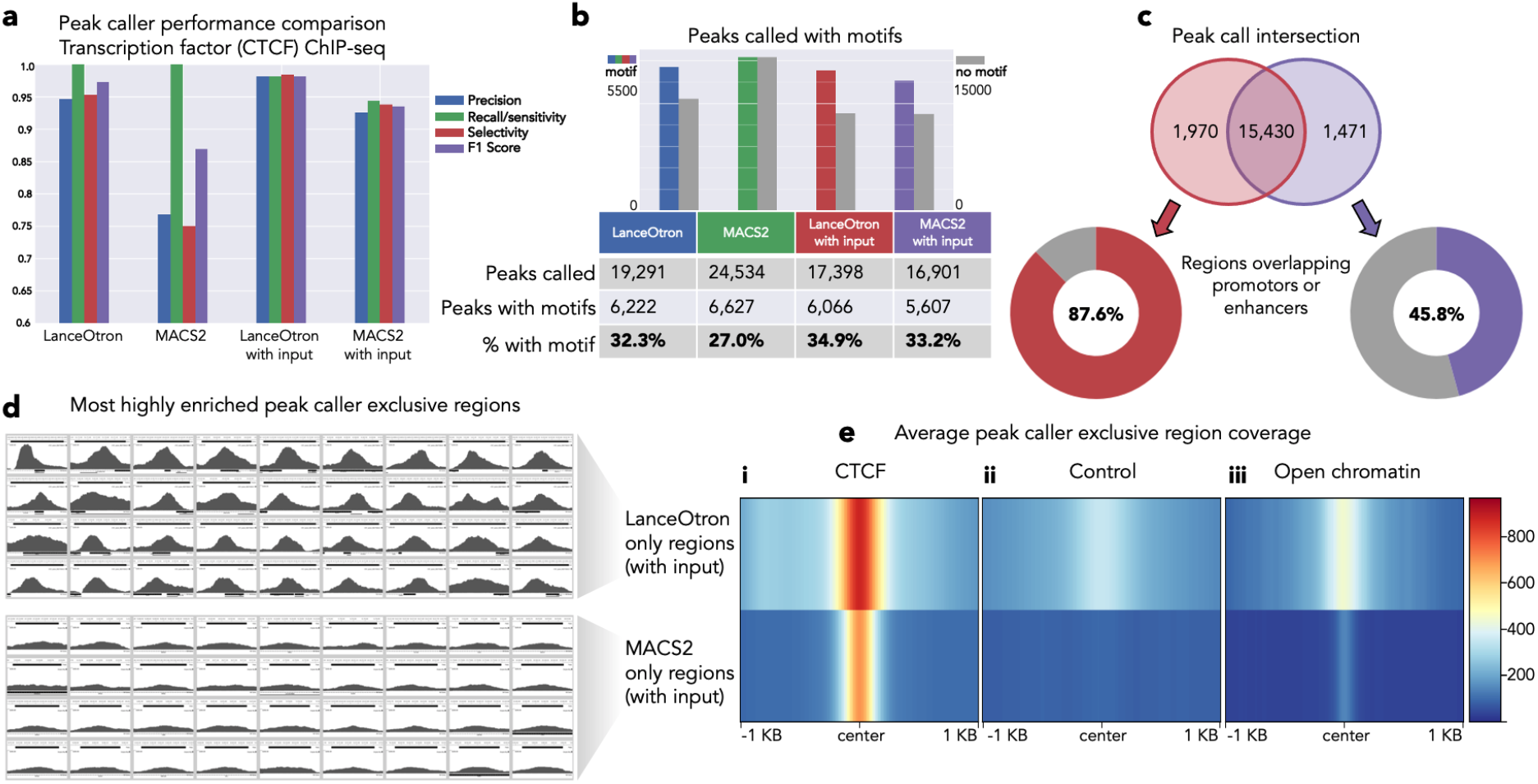
Benchmarking LanceOtran against MACS2 for peak calling transcription factor ChIP-seq. **a**, Model performance metrics using labelled genomic regions of an ENCODE CTCF ChIP-seq dataset. **b**, Comparing the number of motifs contained the in peak calls generated from LanceOtran and MACS2. **c**, Venn diagram of peak calls from LanceOtran and MACS2. Regions which did not intersect were assessed for overlap with promotors enhancers. **d**, Thumbnail images from the most highly enriched regions called exclusively by either LanceOtran (top) or MACS2 (bottom). **e**, Average coverage of the regions called exclusively by either LanceOtran (top) or MACS2 (bottom) for CTCF experimental track, control track, and DNase-seq open chromatin track.

To gain insight into the peak calls genome wide, we performed motif analysis. The number of peaks called were similar between the different methods, though MACS2 without input was slightly higher: LanceOtron, 19,291; MACS2, 24,534; LanceOtron with input, 17,398; MACS2 with input, 16,901. Without input, LanceOtron called fewer peaks with motifs than MACS2 but called fewer peaks in total, resulting in a larger percentage of the overall peak call containing motifs: 32.3% for LanceOtron versus 27.0% for MACS2. When inputs were used, LanceOtron had both a larger count of peaks containing CTCF motifs as well as a larger percentage of the peak call with motifs: 34.9% versus 33.2% (**Fig. 2b**).

We further investigated the differences between LanceOtron with input and MACS2 with input peak calls, finding 1,970 LanceOtron only and 1,471 MACS2 only regions. The transcription factor being tested for in this experiment, CTCF, is often associated with promoters and enhancers^31^, and we found 87.6% of peaks found exclusively with LanceOtron overlapped with promoters or enhancers compared to just 45.8% of MACS2 only peak calls (**Fig. 2c**). When visualizing the top enriched regions called exclusively by each peak caller, LanceOtron’s peaks have strikingly more signal than MACS2 (**Fig. 2d**). Indeed, this trend holds when inspecting the average signal of the exclusive peak calls; MACS2 only regions were found in regions with less surrounding signal, containing peaks which were narrower and with very low enrichment compared to LanceOtron only peaks. It seems that the MACS2-only regions are a sporadic sampling of the numerous peaks close to noise found throughout the genome, however the peaks that MACS2 missed are relatively strongly enriched. These missing peaks are excluded by MACS2 because of the increase in control signal, however some increased signal from the control track is expected when the region is found in areas of open chromatin^6^, which can be seen associated with the LanceOtron only peaks (**Fig. 2e**). Using the outcome of a statistical test as the sole criteria of categorizing genomic regions means striking a balance between calling false positives and false negatives. While the stringent cut-off set by MACS2 helps reduce false positives genome wide, here it does so at the cost of false negatives. Because LanceOtron additionally uses the shape of the peak, the statistical threshold can be relaxed, thus preventing these false negatives without trading them for a plethora of false positives.

#### Histone ChIP-seq

Our histone ChIP-seq datasets were H3K27ac in HAP-1 cells (ENCSR131DVD) and H3K4me3 in MG63 cells (ENCSR579SNM). For H3K27ac, the top sensitivity was achieved with three peak calls: LanceOtron, both with and without input, and MACS2 without input. LanceOtron outperformed MACS2 in the remaining metrics of precision, selectivity, and F1 score. The same performance was achieved both with and without input for the LanceOtron peak calls, highlighting the power of its deep neural network (LanceOtron HAP-1 H3K27ac projects: without input; with input)(**Fig. 3a**). In the H3K4me3 dataset, specificity was equal between LanceOtron and MACS2 with input, and LanceOtron outperformed MACS2 across all peak call types for the remaining metrics (LanceOtron MG63 H3K4me3 projects: without input; with input)(**Fig. 3b**).

**Fig. 3.**
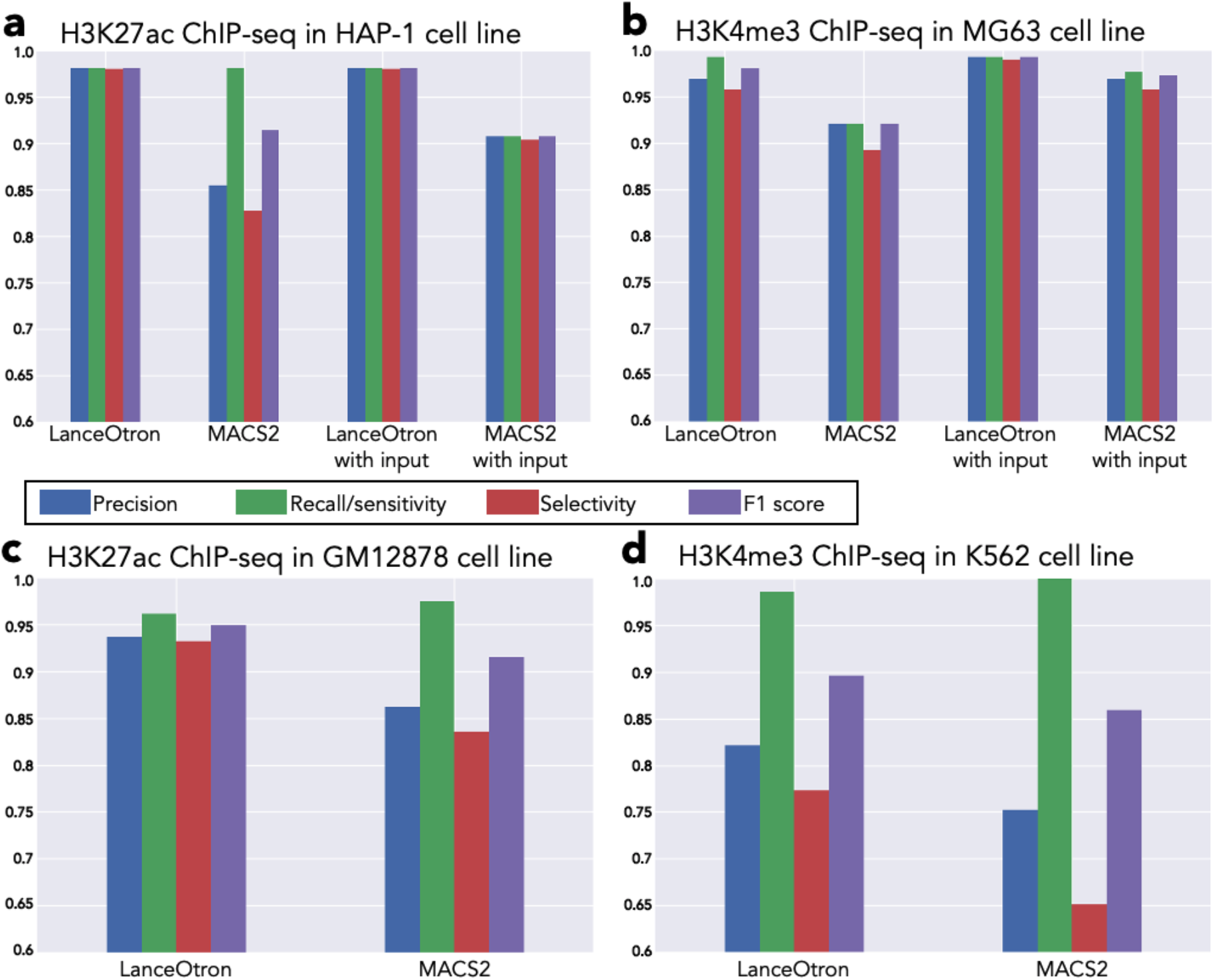
Benchmarking LanceOtron against MACS2 for peak calling histone ChIP-seq. **a**, Model performance metrics using 10 Mb of labelled genomic regions of ENCODE ChIP-seq datasets for H3K27ac in HAP-1 cell line, and **b**, H3K4me3 in MG63 cell line. **c**, ChIP-seq dataset labelled by Oh et al. for H3K27ac in GM12878 cell line and **d**, H3K4me3 in K562 cell line.

We also tested published datasets from Oh et al., who annotated peaks and noise for H3K27ac ChIP-seq in GM12878 cells and H3K4me3 in K562 cells^27^. Performance was generally consistent with our in-house labeled data, and though MACS2 performed slightly better than LanceOtron on sensitivity, LanceOtron outperformed MACS2 on precision, selectivity, and F1 score for both the H3K27ac data (LanceOtron GM12878 H3K27ac project)(**Fig. 3c**) and H3K4me3 data (LanceOtron K562 H3K4me3 project)(**Fig. 3d**).

To further investigate the histone mark ChIP-seq peak calls, we counted the number of transcription start sites (TSSs) overlapping with the peak calls, as TSSs are generally modified with H3K27ac and H3K4me3. Due to the frequency with which TSSs are found in the genome, we restricted the analysis to the top 5,000 peaks called for each peak caller and normalized the regions’ size to 1 kb. This had the added benefit of being resilient to peak caller parameter changes, as the top peaks were unlikely to change based on parameters. For H3K27ac, LanceOtron performance was very similar with and without input, increasing from 2,806 to 2,812 peaks when the input track was included. Both LanceOtron peak calls had more overlap with TSSs than MACS2, which had 2,367 and 2,591 with input. We observed similar results for the H3K4me3 data, with LanceOtron finding 3,472 peaks intersecting TSSs, increasing slightly to 3,501 with input control. MACS2 had better performance without input, though not reaching LanceOtron levels, at 3,335 and decreasing down to 2,589 with input (**Table 1**).

**Table 1.** LanceOtron and MACS2 peak call comparison for transcription start sites (TSSs), and for active regions in open chromatin. Percentages and counts of peaks intersecting TSSs are given for 5,000 regions of LanceOtron and MACS2 peak calls, selected for being most enriched (highest Peak Score or q-value for LanceOtron and MACS2 respectively). Percentages and counts are also shown for oen chromatin peaks found in active areas of the genome.

| Top enriched peaks intersecting TSSs |  |  |
| --- | --- | --- |
|  | LanceOtron | MACS2 |
| % top H3K27ac ChIP-seq in HAP-1 peaks overlapping TSSs (count) | 56.1% (2,806 / 5,000) | 47.3% (2,367 / 5,000) |
| % top H3K4me3 ChIP-seq in MG63 peaks overlapping TSSs (count) | 69.4% (3,472 / 5,000) | 66.7% (3,335 / 5,000) |
| % top ATAC-seq in MCF-7 peaks overlapping TSSs (count) | 44.4% (2,218 / 5,000) | 21.7% (1,086 / 5,000) |
| % top DNase-seq in A549 peaks overlapping TSSs (count) | 43.3% (2,164 / 5,000) | 23.0% (1,151 / 5,000) |
|  | LanceOtron with input | MACS2 with input |
| % top H3K27ac ChIP-seq in HAP-1 peaks overlapping TSSs (count) | 56.2% (2,812 / 5,000) | 51.8% (2,591 / 5,000) |
| % top H3K4me3 ChIP-seq in MG63 peaks overlapping TSSs (count) | 70.0% (3,501 / 5,000) | 51.8% (2,589 / 5,000) |
| GM12878 open chromatin peaks intersecting active regions |  |  |
|  | LanceOtron | MACS2 |
| % ATAC-seq peaks in active regions (count) | 9.2% (5,648 / 60,962) | 7.1% (6,679 / 94,197) |
| % DNase-seq peaks in active regions (count) | 11.4% (2,871 / 25,183) | 7.8% (5,285 / 67,461) |

#### ATAC-seq and DNase-seq

In-house data for ATAC-seq consisted of regions in the MCF-7 cell line from ENCODE (ENCSR422SUG). LanceOtron outperformed MACS2 across all metrics (LanceOtron MCF-7 ATAC-seq project)(**Fig. 4a**). Results were similar for our in-house DNase-seq data in the A549 cell line from ENCODE (ENCSR000ELW). MACS2 outperformed LanceOtron for recall/sensitivity but had a very high false positive rate. Consequently, LanceOtron outperformed MACS2 on precision, sensitivity, and F1 score (LanceOtron A549 DNase-seq project)(**Fig. 4b**). As with the histone datasets, we also intersected the top 5,000 peaks, normalized to 1 kb, with TSSs. LanceOtron’s top peaks had nearly double the number of intersections with TSSs compared with MACS2 for DNase-seq (2,164 versus 1,151) and over double for ATAC-seq peaks (2,218 versus 1,086) (**Table 1**).

**Fig. 4.**
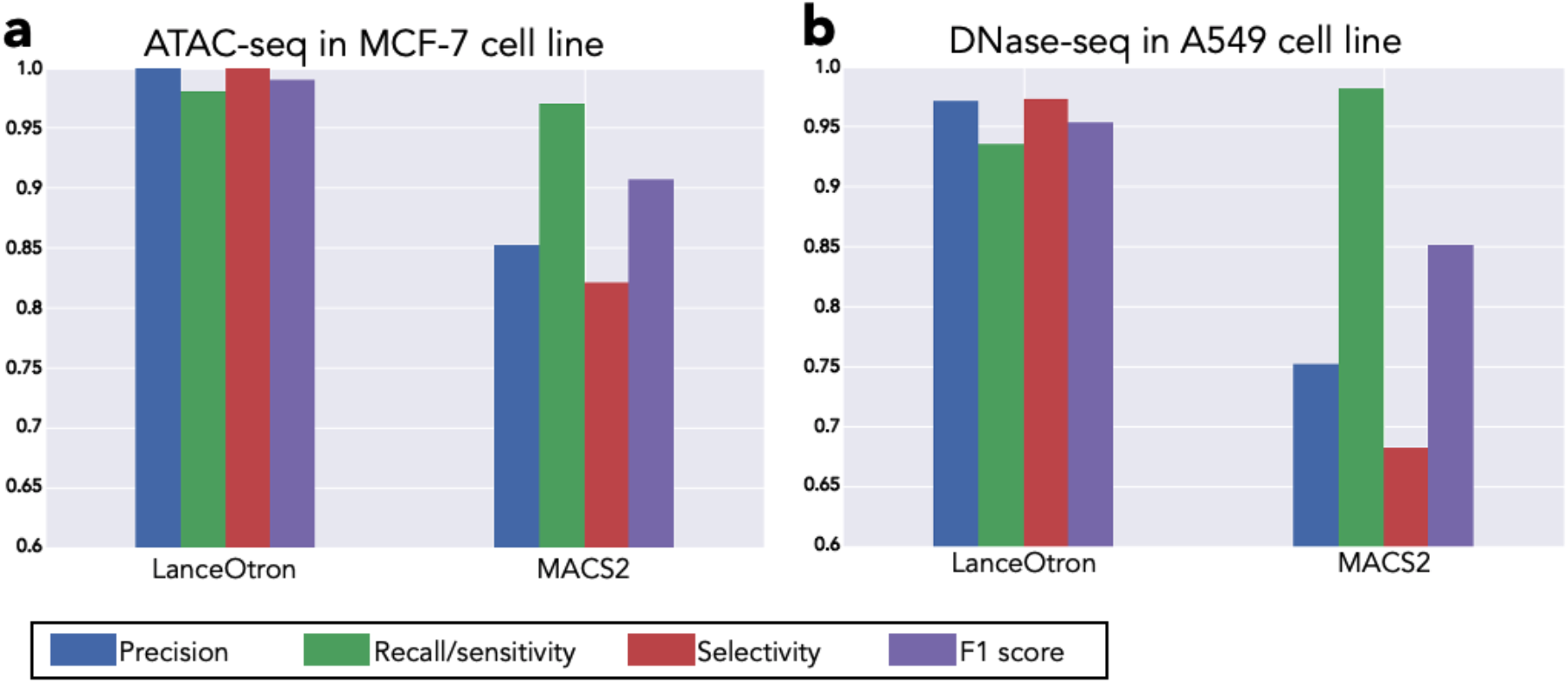
Benchamrking LanceOtron against MACS2 for calling open chromatin. **a**, Model perfromance metrics using labelled genomic regions of an ENCODE ATAC-seq dataset in MCF-7 cell line and **b**, DNase-seq in A549 cell line.

We also compared peak calling performance on GM12878 cells for ATAC-seq (ENCFF576DMC) and DNase-seq (ENCSR000EMT). Here we used published annotations from Tarbell and Liu^32^, whereby they defined active areas of the genome using enhancer and promoter data with the software GenoSTAN^33^. The number of peaks called for these datasets were considerably different from each peak caller. For ATAC-seq, 60,962 peaks were called using LanceOtron and 94,197 peaks for MACS2; DNase-seq, 25,183 peaks were called using LanceOtron and 67,461 peaks for MACS2. For both ATAC-seq and DNase-seq the raw counts of peaks found in active areas were higher with MACS2, however as a percentage of the total peak call, MACS2 was lower than LanceOtron for both experiment types (LanceOtron GM12878 projects: ATAC-seq; DNase-seq)(**Table 1**).

## Discussion

LanceOtron is a deep learning based peak caller for genomic signal analysis, with a full user-friendly interface designed for interrogation of large datasets. Here it outperformed the current gold standard algorithm, MACS2, in each of our experiments. LanceOtron’s CNN, trained on open chromatin, transcription factor and chromatin modification data, learns the shape of the signal and uses this in combination with enrichment calculations to identify biologically relevant regions. Traditional peak callers return only those regions which cross a high statistical threshold. When using LanceOtron’s candidate peak calling algorithm however, all enriched regions above a relatively low threshold are returned, along with their associated peak scores, p-values, heights, widths, and other properties. This makes LanceOtron akin to an automated annotation tool, returning a greater breadth of data about the experiment. It’s function as a peak caller is realized using LanceOtron’s comprehensive data visualization, filtering and data handling to generate output data sets with defined characteristics.

Benchmarking the transcription factor CTCF ChIP-seq data showed that peaks uniquely identified by LanceOtron were enriched for enhancers or promoters, as would be expected based on the biology of the transcription factor being analyzed. In our testing some real peaks appeared to be absent from the MACS2 dataset. Inspection of the DNase-seq track made it clear that many of the regions missed by MACS2 were in regions of open chromatin. These areas sonicate more readily^6^, and are known to have increased signal in input tracks, however this increase in control signal did not preclude these regions from being recognized using LanceOtron as it did for MACS2. The loss of these regions in the MACS2 analysis is likely due to a combination of its reliance on the signal in the input tracks and its high p-value threshold to better reduce false positives genome wide, but at the cost of sensitivity in active regions of the genome. LanceOtron peaks were shown to be enriched for the CTCF binding motif more often than MACS2, providing biological evidence that the differences in peaks called by LanceOtron are actually improvements over traditional analysis. Though outperformed by LanceOtron, MACS2 actually achieved its second-best overall performance score on the CTCF dataset (F1 score, first best on H3K4me3). This is perhaps not surprising as MACS2 was designed for peak calling transcription factor binding experiments. While the narrow binding pattern of H3K4me3 closely resembles the distribution of read coverage seen in transcription factor binding, MACS2 performance suffered when the genomic signal deviated from this pattern, especially true in ATAC-seq and DNase-seq. When no input track was available, MACS2 overall performance further declined. This is in contrast to LanceOtron, whereby performance was only slightly lower without an input track, and even outperformed MACS2 with input for overall F1 score on every test where this comparison was available (**Supplementary Table 2**).

LanceOtron’s dual focus of deep learning on big data and generation of rich interactive visualizations are each computationally expensive in their own right. Yet despite this, the average time to perform a peak call on the 13 datasets benchmarked here, plus automatically generate the interactive charts and genome browser, was just over an hour to run *within a web browser* (mean time 67 minutes, standard deviation 11 minutes). The speed that LanceOtron can carry out analysis, requiring only a basic bigwig track and using a web interface, also means that it can form part of the review process when a manuscript is under consideration. While often a session of data is provided during review, this is seldom utilized due to time constraints and the necessity of accessing high performance computing facilities. LanceOtron remedies this, providing a convenient outlet for group leaders, bench biologists, and bioinformaticians alike to visualize and assess from internal or external sources. In addition, peak calls made with LanceOtron can easily be made public for assessment by reviewers and colleagues directly, as they have been here. Improving access to the analysis process is beneficial to the larger molecular biology community and helps to address the growing concern of reproducibility in science.

LanceOtron had a comprehensive development process during which over 100 unique users tested the tool, with over 30 users creating 10 projects or more. We have learned how labs around the world analyze their chromatin profiling assays, and we designed our workflow around this experience. A strength of using supervised machine learning approaches is that analysis can improve as more training data is added to the model; as our user base grows, we can refine our peak calls even further. Our focus to date has been on the most commonly used experiments where we believed there was the greatest potential for improvement. However, unlike hardwired statistical algorithms, CNN-based algorithms can easily be trained to deal with new signal types and distributions not covered in the original training sets. The same architecture can potentially be used to learn different types of genomics data, for example CAGE transcription start site signals or methylomics which are currently challenging to extract signal from noise; exemplifying this, LanceOtron has even been adapted for analysing base pair resolution chromosome conformation capture^34^.

In summary, LanceOtron is a powerful peak caller and analysis tool for ATAC-seq, ChIP-seq, and DNase-seq. Across a range of different datasets and data types, LanceOtron outperformed the industry-standard MACS2. It is designed to accommodate current workflows as a visualization, annotation, filtering and peak calling tool, leveraging a powerful deep learning neural network to use peak shape information alongside enrichment data.

## Methods

### Deep learning model

#### Training data

The data used to train the neural network was obtained from ENCODE. To generate a complete list of experiments which met our specifications we used ENCODE’s REST API (scripts and outputs available on GitHub). We filtered the results to samples which were “released” status at the time of search inquiry and aligned to human reference genome hg38 as BAM files; for H3K27ac, H3K4me3, and transcription factor ChIP-seq experiments, the availability of a corresponding control track was also required. While infrequent, samples were excluded if ENCODE metadata did not include information on single-end versus paired-end sequencing. The number of samples meeting these criteria was 3,902 (74 ATAC, 911 DNase, 305 H2K27ac, 463 H3K4me3, 2,149 transcription factor samples). We sampled 10 paired-end datasets for each category at random from each experiment type, except in H3K4me3 experiments where only 6 samples available were paired-end, and so 4 single end experiments were included. This resulted in 38 unique biosample types, 9 unique transcription factor ChIP-seq targets plus 2 histone ChIP-seq targets (**Table 2**).

Each BAM file was downloaded directly from ENCODE, along with the corresponding control BAMs for H3K27ac, H3K4me3, and transcription factor ChIP-seq experiments. If multiple replicates of the control experiments existed, only the first listed in ENCODE’s database was used for analysis. BAM files were sorted and indexed using Samtools^35^ 1.3 (samtools sort filename.bam and samtools index filename.bam.sorted commands respectively). Bigwig file coverage maps were created from the BAM files using deepTools^36^ version 3.0.1 commands: `bamCoverage --bam filename.bam.sorted -o filename.bw --extendReads -bs 1 --normalizeUsing RPKM` for paired-end sequenced experiments. For single-end sequenced experiments the average fragment length was obtained from ENCODE and used with the --extendReads flag, making the command: `bamCoverage --bam filename.bam.sorted -o filename.bw --extendReads averageFragmentLength -bs 1 --normalizeUsing RPKM`.

Putative peak calls were carried out on all datasets, followed by classification as either peak or noise based on visual inspection. Coordinates for the regions being assessed were determined three ways. The MACS2 peak caller was used on default settings, `macs2 callpeak -t filename.bam.sorted -c control_filename.bam.sorted -n sample_label -f BAM -g hs -B -q 0.01` for H3K27ac, H3K4me3, and transcription factor ChIP-seq datasets. For ATAC-seq and DNase-seq, which lack control tracks, the following command was used: `macs2 callpeak -t filename.bam.sorted -n sample_label -f BAM -g hs -B -q 0.01`. The second and third peak call methods focused on labeling regions based on their fold enrichment compared to the mean signal. Coverage maps of sequenced reads were first smoothed by applying a rolling average of a given window size. If this smoothed signal was greater than the mean multiplied by a fold enrichment threshold, the coordinate was marked as enriched; adjacent enriched regions were then merged. Methods two and three used five smoothing windows at different base pair (bp) resolutions (100 bp, 200 bp, 400 bp, 800 bp, 1600 bp) as well as five different enrichment thresholds (1, 2, 4, 8, 16). Method two compared the smoothed signal to the mean of chromosome-wide signal multiplied by fold enrichment. Method three was similar except the smoothed signal was compared to either the mean of the chromosome, surrounding 5 kb, or surrounding 10 kb, whichever value was highest (i.e. `max[chromosome mean, 5 kb mean, 10 kb mean]`) multiplied by fold enrichment.

From each dataset a 1 Mb continuous region was selected at random for each chromosome for autosomes and sex chromosomes only. If the start of the randomly selected region was near the end of the chromosome, the area considered was from that point to the chromosome end, then from the chromosome start extending out until a full 1 Mb was covered. Peaks called from all 3 methods which started within the random region were made available for labeling. For both of the mean-based methods, a peak call was made for each permutation of the smoothing window and enrichment threshold parameters, and all 25 calls were combined - this meant the presence of multiple overlapping candidate peaks in some cases. A python implementation of BEDTools^37^ (pybedtools) was used to find overlapping peaks, and only one selected at random was considered for visual inspection.

Only candidate regions which were obviously peaks or noise were labeled as such. Visual inspection was carried out using MLV^15^, with control tracks overlaid when available. Regions were inspected one at a time, until 100 verified peaks were found for the dataset or all of the regions were assessed. Entire 1 Mb regions were assessed (no early stopping), with the order of chromosomes randomized. A total of 736,753 regions were labeled this way (5,016 peaks and 731,737 noise regions) covering 499 Mb.

Additional labels were generated using an algorithm. First the raw signal was smoothed by calculating the rolling mean for the surrounding 400 bp, and any coordinate where the signal was 4-fold*mean-chromosome-signal was marked as enriched. Adjacent enriched regions were combined, and if the size was between 50 bp and 2 kb it was considered a candidate peak. Regions smaller than 50 bp were discarded, and regions above 2 kb were recursively re-evaluated at a 1-fold higher threshold until the region size was between 50 bp and 2 kb, or the region was greater than 20-fold enriched. If these candidate peaks intersected with the previously labeled peaks, these regions were then also labeled peaks, resulting in an additional 3,447 labels for a total of 8,463 peaks (ATAC-seq: 1,926; DNase-seq: 2,097; H3K27ac ChIP-seq: 1,651; H3K4me3 ChIP-seq: 1,806; transcription factor ChIP-seq: 983). Noise regions were down sampled with prioritization given to regions with the highest signal. All noise regions with a max height in the 25th percentile or greater were included (3,658), and equal numbers below the 25th percentile were randomly sampled.

These labeled data were used for training the first phase of the model. Afterwards we scored the training data with this model to identify any mislabeled data or model misclassifications; from this process 24 peaks and 1,187 noise regions were added to the dataset. Ultimately 16,990 regions were used for training: 8,503 noise regions plus 8,463 peaks.

#### Wide and deep convolutional neural network to learn shape and enrichment of regions

LanceOtron’s machine learning architecture is a type of wide and deep neural network^38^, combining enrichment values, logistic regression, and a CNN. The logistic regression model takes as inputs the enrichment values, while the CNN uses the 2 kb of signal centered on the region of interest. The outputs of these two models, along with the 11 enrichment values, are input into a multilayer perceptron, which outputs a peak quality metric (called Peak Score) with values ranging from 0 to 1.

The 11 enrichment values consisted of Poisson-based p-values, using maximum height and average signal, calculated from 10kb to 100kb regions in 10kb increments as well as chromosome-wide enrichment. While this is an internal model parameter, and not used for significance thresholding, we opted to use the p-value because of the increased interpretability, though numerous enrichment metrics could have been used to yield similar results. These p-values are also returned to the user as an additional calculated measurement; indeed the results of a traditional statistical peak caller could be mimicked by simply using p-values as the sole filtering metric.

The logistic regression model was trained separately with the same training data, and all coefficients and model parameters saved. The wide and deep model was then trained with the logistic regression component locked, and with loss distributed 70:30 to wide-and-deep-output:CNN-only-output. By penalizing the model on the CNN separately, it actively encouraged predictions from the 2 kb of signal, i.e., the shape of the peak, to be accurate in absence of enrichment information.

To determine the optimal structure and hyperparameters, a brute force method of building many models with different configurations was carried out. In total 5,000 models were trained and tested using the python package Keras Tuner, though performance was robust across a range of configurations. Model performance was assessed by measuring the number of correctly predicted classifications of enriched regions from data unseen to the model. The top 10 performing models were then subjected to 5-fold cross validation, and the architecture from the top performer was used.

### Candidate peak selection

To optimize resources, candidate peaks are selected for their enrichment above the mean chromosome signal, whereby the signal is extracted and passed to LanceOtron’s neural network. We developed an algorithm which acts as a loose filter, allowing even modestly enriched regions through, which also centers the area around the highest signal and improves model performance. First the raw signal is smoothed by calculating the rolling mean for five different window sizes, 100 bp, 200 bp, 400 bp, 800 bp, and 1600 bp. Next any coordinate where the signal is greater than fold*mean-chromosome-signal (across 5 different fold enrichments: 2, 4, 8, 16, and 32) is marked as enriched. Each permutation of the rolling mean window size and fold threshold is considered a different definition of enrichment. The number of enrichments is tracked at each coordinate, forming a genome wide map, and regions with 5 or more concurring definitions of enrichment are further evaluated. If the region’s size is between 50 bp and 2 kb it is considered a candidate peak. Regions smaller than 50 bp are discarded, and regions above 2 kb are recursively increased by an additional required enrichment definition until the region size is between 50 bp and 2 kb, or the region is considered enriched under all 25 definitions.

### Calculating p-value from an input control track

A standard p-value assessment based on the Poisson distribution is performed when using LanceOtron’s Find and Score Peaks with Inputs module, which can be used in conjunction with the peak quality metric output from LanceOtron’s deep learning model. The mean signal expected from background, *λ*, is determined using either the mean signal in the input control track (*λ*_input_) or the mean signal in the input control track plus 1 kb (*λ*_1kb_), whichever is more stringent. P-values are then computed using the average count of overlapping reads (N_ave_) within the given candidate region.

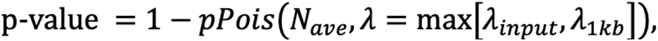

where *pPois* is the Poisson cumulative distribution function:

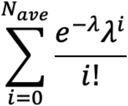

### Peak caller benchmarking

#### Labeling testing data and calculating model performance

Testing datasets were also obtained from ENCODE but were not used in LanceOtron’s training data (**Supplementary Table 1**). Each track was downloaded as a BAM file and converted to bigwig using the same deepTools commands given above for in training data preparation. Chromosomes were shuffled (mitochondrial and alternative mapping chromosomes were excluded), and 1 Mb was labeled for peaks or noise; regions which were not clearly either were excluded. For CTCF, H3K27ac, and H3K4me3 ChIP-seq datasets, 10 chromosomes each were labeled in this manner for a total of 122 labels (55 positive peaks and 67 noise regions), 101 labels (45 positive peaks and 56 noise regions), and 224 labels (129 positive peaks and 95 noise regions) respectively. For ATAC-seq and DNase-seq, 3 chromosomes each were labeled, resulting in 196 ATAC labels (101 positive peaks and 95 noise regions) and 224 DNase labels (114 positive peaks and 110 noise regions). True positives, false positives, true negatives, and false negatives were determined by intersecting peak calls from LanceOtron and MACS2 with these labeled data using BEDTools. True positives were found by using the command `bedtools intersect -a peak_call.bed -b labeled_peaks.bed -u -wa`. False negatives used `bedtools intersect -a peak_call.bed -b labeled_peaks.bed -v -wa`. True negatives used the command `bedtools intersect -a peak_call.bed -b labeled_noise.bed -v -wa`, while false positives used `bedtools intersect -a peak_call.bed -b labeled_noise.bed -u -wa`.

For peaks which were exclusively found with LanceOtron or MACS2, BEDTools was also used to find intersections which occurred at promoter or enhancer regions, as well as TSSs. The bed files listing the coordinates of the promoters or enhancers were from GenoSTAN^33^, and for TSSs we used RefTSS^39^.

The heat map of the coverage was made using the deeptools command: computeMatrix reference-point -S CTCF_spleen_ENCFF656CCY.bw -R CTCF-spleen_LoT-only-peaks.bed CTCF-spleen_MACS2-only-peaks.bed --referencePoint center -a 1000 -b 1000 -out CTCF-spleen_LoT-and-MACS2_matrix.tab.gz Followed by the command: plotProfile -m CTCF-spleen_LoT-and-MACS2_matrix.tab.gz -out CTCF-spleen_LoT-and-MACS2.png --samplesLabel “Peak caller exclusive regions” --regionsLabel “LanceOtron only” “MACS2 only” --plotType=heatmap.

#### Motif analysis

A custom motif matching script was written to match CTCF sites using a simple Python regex function. The motif position weight matrix (PWM) was downloaded from JASPAR^40^ and the genomic coordinates matching the motif (and reverse complement) were recorded as a bed file. The matching sequence had to be the same length, with all nucleotides present at 75% or higher in the PWM as exact matches. With the bed file of the motif coordinates made, we once again employed BEDTools to find intersections with the peak calls.

## Code availability

The LanceOtron webtool peak caller can be used at https://LanceOtron.molbiol.ox.ac.uk/ with the source code and command line algorithm implementation found at https://github.com/LHentges/LanceOtron.

## Acknowledgments

The authors would like to thank Dr. Simon J. McGowan for web design insight, project discussions and proofreading, as well as Dr. Jon Kerry and Dr. Dominic Waithe for their pilot research. We also want to acknowledge the many beta testers and users during the software development, in particular efforts from the Hughes Lab, AVI Group, Higgs Group, Milne Group, and the Centre for Computational Biology at the MRC Weatherall Institute of Molecular Medicine, University of Oxford. This work was also supported by the National Institutes of Health (USA) (R24DK106766 to J.R.H.), the Medical Research Council (MC_UU_12025 to S.T. and MC_UU_00016/14 to J.R.H.) and a Wellcome Trust Strategic Award (106130/Z/14/Z to J.R.H.).

## Author contributions

J.R.H. and S.T. designed the project and directed the research. L.D.H. built the candidate peak calling algorithm, labeled the training data, coded, trained, and tested the deep learning model, and created the command line tool. M.J.S. designed and coded the graphical user interface as well as the interactive visualization tools and built the website. D.J.D., J.R.H., and S.T. tested the software and suggested new features.

## Supplementary Materials

### Functionality of LanceOtron’s user interface

LanceOtron features a rich graphical user interface, accessible using any web browser, and allows peak calls to be made without the use of the command line. Using the web tool to perform a peak call is demonstrated in **supplementary video 1**: https://youtu.be/k8GrIp55vDg. Furthermore, exploring and filtering data is also easily carried out with the graphical interface, demonstrated in **supplementary video 2**: https://youtu.be/M5ox8XI-U4Q.

**Supplementary Table 1.** ENCODE datasets used for training data and testing data

| Training Data |  |  |  |  |  |
| --- | --- | --- | --- | --- | --- |
| Experiment |  |  | ENCODE ID numbers |  |  |
| Assay | Target | Tissue | Experiment | BAM file | Control BAM |
| ATAC-seq | Open chromatin | Breast epithelium | ENCSR955JSO | ENCFF656OYT |  |
| ATAC-seq | Open chromatin | Tibial artery | ENCSR630REB | ENCFF168OTV |  |
| ATAC-seq | Open chromatin | Foreskin keratinocyte | ENCSR290YMN | ENCFF799HAR |  |
| ATAC-seq | Open chromatin | Adrenal gland | ENCSR113MBR | ENCFF436NOT |  |
| ATAC-seq | Open chromatin | Foreskin keratinocyte | ENCSR158XTU | ENCFF784DSJ |  |
| ATAC-seq | Open chromatin | Foreskin keratinocyte | ENCSR677MJF | ENCFF764CQI |  |
| ATAC-seq | Open chromatin | Transverse colon | ENCSR668VCT | ENCFF377DAO |  |
| ATAC-seq | Open chromatin | Sigmoid colon | ENCSR548QCP | ENCFF482HAC |  |
| ATAC-seq | Open chromatin | Tibial nerve | ENCSR831KAH | ENCFF277DNH |  |
| ATAC-seq | Open chromatin | Thyroid gland | ENCFF710ELD | ENCSR474XFV |  |
| ChIP-seq | H3K27ac | RWPE1 | ENCSR203KEU | ENCFF708CBX | ENCFF939LTT |
| ChIP-seq | H3K27ac | SKNSH | ENCSR564IGJ | ENCFF380OTV | ENCFF959FMO |
| ChIP-seq | H3K27ac | Bipolar neuron | ENCSR905TYC | ENCFF751YAL | ENCFF687LIL |
| ChIP-seq | H3K27ac | GM23338 | ENCSR729ENO | ENCFF403VXK | ENCFF754UFV |
| ChIP-seq | H3K27ac | C42B | ENCSR279KIX | ENCFF913EZV | ENCFF980IJT |
| ChIP-seq | H3K27ac | 22Rv1 | ENCSR391NPE | ENCFF025ZEN | ENCFF769UET |
| ChIP-seq | H3K27ac | Foreskin keratinocyte | ENCSR709ABP | ENCFF085FAH | ENCFF178GZR |
| ChIP-seq | H3K27ac | Foreskin keratinocyte | ENCSR709ABP | ENCFF776HMQ | ENCFF178GZR |
| ChIP-seq | H3K27ac | Epithelial cell of prostate | ENCSR910PDW | ENCFF382XYO | ENCFF213AZI |
| ChIP-seq | H3K27ac | RWPE2 | ENCSR987PNT | ENCFF245ORL | ENCFF169DGZ |
| ChIP-seq | H3K4me3 | SKNSH | ENCSR975GZA | ENCFF027SGQ | ENCFF959FMO |
| ChIP-seq | H3K4me3 | SKNSH | ENCSR975GZA | ENCFF245RXP | ENCFF959FMO |
| ChIP-seq | H3K4me3 | NCIH929 | ENCSR082NQB | ENCFF417RNS | ENCFF446RUP |
| ChIP-seq | H3K4me3 | NCIH929 | ENCSR082NQB | ENCFF067LLV | ENCFF446RUP |
| ChIP-seq | H3K4me3 | Bipolar neuron | ENCSR849YFO | ENCFF096QTT | ENCFF687LIL |
| ChIP-seq | H3K4me3 | Bipolar neuron | ENCSR849YFO | ENCFF950QWN | ENCFF687LIL |
| ChIP-seq | H3K4me3 | Muscle of leg | ENCSR128QKM | ENCFF552OGD | ENCFF622XBJ |
| ChIP-seq | H3K4me3 | Heart right ventricle | ENCSR107RDP | ENCFF897OOT | ENCFF246SXV |
| ChIP-seq | H3K4me3 | Gastrocnemius medialis | ENCSR098QLN | ENCFF310NMI | ENCFF587DDD |
| ChIP-seq | H3K4me3 | OCILY3 | ENCSR548PZS | ENCFF816RLY | ENCFF691EEI |
| ChIP-seq | NR2C1 | GM12878 | ENCSR784VIQ | ENCFF785FLS | ENCFF322NTO |
| ChIP-seq | EP300 | Ovary | ENCSR696LQU | ENCFF405UYE | ENCFF271JKY |
| ChIP-seq | NFXL1 | GM12878 | ENCSR746XEG | ENCFF673BXM | ENCFF322NTO |
| ChIP-seq | MXI1 | Neural cell | ENCSR934NHU | ENCFF260PNL | ENCFF056HWK |
| ChIP-seq | ZNF318 | K562 | ENCSR334HSW | ENCFF373YTD | ENCFF790TAN |
| ChIP-seq | CREB1 | HepG2 | ENCSR112ALD | ENCFF011HOS | ENCFF950AXC |
| ChIP-seq | CTCF | RWPE1 | ENCSR303GFI | ENCFF204KRO | ENCFF290UZX |
| ChIP-seq | RFX1 | MCF7 | ENCSR788XNX | ENCFF804LEF | ENCFF426RDP |
| ChIP-seq | CTCF | Ascending aorta | ENCSR960MDF | ENCFF353ZVY | ENCFF023NJF |
| ChIP-seq | E4F1 | K562 | ENCSR731LHZ | ENCFF978NVP | ENCFF910IKB |
| DNase-seq | Open chromatin | Left arm bone | ENCSR976XOY | ENCFF205JXZ |  |
| DNase-seq | Open chromatin | A673 | ENCSR346JWH | ENCFF348KWA |  |
| DNase-seq | Open chromatin | T-helper 1 cell | ENCSR000EQC | ENCFF425YMJ |  |
| DNase-seq | Open chromatin | Retina | ENCSR820ICX | ENCFF441YDL |  |
| DNase-seq | Open chromatin | Uterus | ENCSR129BZE | ENCFF759POB |  |
| DNase-seq | Open chromatin | NAMALWA | ENCSR301OGM | ENCFF554YJG |  |
| DNase-seq | Open chromatin | SKMEL5 | ENCSR000FEK | ENCFF844BZM |  |
| DNase-seq | Open chromatin | ELF1 | ENCSR678ILN | ENCFF433CFI |  |
| DNase-seq | Open chromatin | Myocyte | ENCSR000EPD | ENCFF042QTI |  |
| DNase-seq | Open chromatin | Pancreas | ENCSR828FVZ | ENCFF984FKS |  |
| **Testing Data** |  |  |  |  |  |
| **Experiment** |  |  | **ENCODE ID numbers** |  |  |
| **Assay** | **Target** | **Tissue** | **Experiment** | **BAM file** | **Control BAM** |
| ATAC-seq | Open chromatin | MCF-7 | ENCSR422SUG | ENCFF346MIJ |  |
| ChIP-seq | CTCF | Spleen | ENCSR692ILH | ENCFF903NKV | ENCFF376BTL |
| ChIP-seq | H3K27ac | HAP-1 | ENCSR131DVD | ENCFF742SZS | ENCFF247DSQ |
| ChIP-seq | H3K4me3 | MG63 | ENCSR579SNM | ENCFF996ZSR | ENCFF381RWF |
| DNase-seq | Open chromatin | A549 | ENCSR000ELW | ENCFF410CDT |  |

**Supplementary Table 2.** numerical listing of performance benchmarks for all datasets

| **CTCF ChIP-seq in spleen** |  |  |  |  |
| --- | --- | --- | --- | --- |
|  | **LanceOtron** | **MACS2** | **LanceOtron with input** | **MACS2 with input** |
| Precision | 0.946 | 0.768 | **0.981** | 0.926 |
| Recall/sensitivity | **1.000** | **1.000** | 0.981 | 0.943 |
| Selectivity | 0.953 | 0.750 | **0.984** | 0.938 |
| F1 score | 0.972 | 0.869 | **0.981** | 0.935 |
| **H3K27ac ChIP-seq in HAP-1** |  |  |  |  |
|  | **LanceOtron** | **MACS2** | **LanceOtron with input** | **MACS2 with input** |
| Precision | **0.981** | 0.854 | **0.981** | 0.907 |
| Recall/sensitivity | **0.981** | **0.981** | **0.981** | 0.907 |
| Selectivity | **0.981** | 0.827 | **0.981** | 0.904 |
| F1 score | **0.981** | 0.914 | **0.981** | 0.907 |

| H3K27ac ChIP-seq in GM12878 |  |  |  |  |
| --- | --- | --- | --- | --- |
|  | LanceOtron |  | MACS2 |  |
| Precision | 0.937 |  | 0.862 |  |
| Recall/sensitivity | 0.961 |  | 0.974 |  |
| Selectivity | 0.932 |  | 0.836 |  |
| F1 score | 0.949 |  | 0.915 |  |
| H3K4me3 ChIP-seq in MG63 |  |  |  |  |
|  | LanceOtron | MACS2 | LanceOtron with input | MACS2 with input |
| Precision | 0.969 | 0.920 | 0.992 | 0.968 |
| Recall/sensitivity | 0.992 | 0.920 | 0.992 | 0.976 |
| Selectivity | 0.957 | 0.892 | 0.989 | 0.957 |
| F1 score | 0.980 | 0.920 | 0.992 | 0.972 |
| H3K4me3 ChIP-seq in K562 |  |  |  |  |
|  | LanceOtron |  | MACS2 |  |
| Precision | 0.821 |  | 0.753 |  |
| Recall/sensitivity | 0.986 |  | 1.000 |  |
| Selectivity | 0.773 |  | 0.652 |  |
| F1 score | 0.896 |  | 0.859 |  |
| ATAC-seq in MCF-7 |  |  |  |  |
|  | LanceOtron |  | MACS2 |  |
| Precision | 1.000 |  | 0.852 |  |
| Recall/sensitivity | 0.980 |  | 0.970 |  |
| Selectivity | 1.000 |  | 0.821 |  |
| F1 score | 0.990 |  | 0.907 |  |
| DNase-seq in A549 |  |  |  |  |
|  | LanceOtron |  | MACS2 |  |
| Precision | 0.971 |  | 0.752 |  |
| Recall/sensitivity | 0.935 |  | 0.981 |  |
| Selectivity | 0.973 |  | 0.681 |  |
| F1 score | 0.953 |  | 0.851 |  |

